# CRISPR/Cas9-mediated elimination of the *LMNA* c.745C>G pathogenic mutation enhances survival and cardiac function in *LMNA*-associated congenital muscular dystrophy

**DOI:** 10.1101/2025.02.13.638060

**Authors:** Déborah Gómez-Domínguez, Carolina Epifano, Iván Hernández, Borja Vilaplana-Martí, Sergi Cesar, Antonio de Molina-Iracheta, Miguel Sena-Esteves, Georgia Sarquella-Brugada, Ignacio Pérez de Castro

## Abstract

*LMNA*-associated congenital muscular dystrophy is a currently incurable rare genetic disorder characterized by early-onset muscle weakness, dilated cardiomyopathy and respiratory failure, resulting from mutations in the *LMNA* gene. In this study, we assessed the potential of a CRISPR-mediated strategy to eliminate the mutant allele *Lmna* c.745C>T, p.R249W using a mutation specific guide (sg745T). Results from R249W-mutation-carrying cellular models showed specific activity of the Cas9/sg745T complex towards the mutant allele. This property varied depending on the concentration of CRISPR components, with a loss of specificity observed with increased dosage. We tested this strategy *in vivo* using adeno-associated virus delivery in *Lmna^R249W^* mice. Despite being associated with a modest CRISPR activity, this therapeutic approach resulted in a 10% increase in the survival of R249W homozygous mice. Interestingly, a similar CRISPR activity improved the cardiac pathology developed by *Lmna^+/R249W^* animals, significantly extending their median survival. These results represent the first therapeutic validation of a CRISPR/Cas9-mediated gene editing strategy for the treatment of *LMNA*-associated congenital muscular dystrophy.

## INTRODUCTION

*LMNA*-related congenital muscular dystrophy (L-CMD; ORPHA:157973) is an autosomal dominant myopathy inherited genetically, with an extremely rare overall prevalence of less than 1 in 1,000,000. L-CMD patients display severe clinical manifestations of skeletal muscle laminopathies, characterized by an early onset and a rapid progression ^1^. The disease is characterized by motor development delay due to significant skeletal muscle weakness observed in the first months of life or even during the fetal period. Symptoms vary in severity and include elevated creatine kinase levels ^2^, dropped head syndrome ^3^, joint contractures ^4^, and cardiac and respiratory complications over time that lead to sudden death ^5^.

L-CMD is one of the fifteen rare diseases designated as laminopathies, which are associated with abnormalities in the nuclear lamina and are caused by mutations in the main components of this sub-cellular structure ^6^. Most L-CMD patients have mutations located in exons 1, 4, 6, and 7 of the *LMNA* gene, which coincide with the coils of the head and central domains of the protein and the immunoglobulin domain of the tail ^5,7^. The most frequent variant associated with L-CMD is the c.745C>T missense mutation, located in exon 4, that results in an arginine to tryptophan substitution at position 249 (p.R249W) ^5,8,9^. To study L-CMD underlying mechanisms and test therapeutic strategies, we have developed and characterized a *Lmna^R249W^* mouse model (manuscript in preparation). *Lmna^R249W/R249W^* mice show severe growth delay leading to premature death at around 50 days of age. On the other hand, *Lmna^+/R249W^* mice recapitulate the cardiac abnormalities developed by L-CMD patients, their genetic equivalents, consisting in progressive dilated cardiomyopathy with left ventricular dilation and reduced cardiac function leading to sudden death.

Currently, there is no cure for L-CMD, and treatment focuses on symptom management. Palliative approaches include exercise programs, mechanical aids, surgical intervention, and monitoring of cardiac and respiratory functions ^10,11^. The disease, despite improvements in life expectancy, remains incurable, emphasizing the need for development of effective therapies. Several pre-clinical studies have assessed the therapeutic potential of different strategies to treat laminopathies. Small molecules drugs targeting the MAPK and mTOR pathways as well as NAT10 have been tested for treatment of laminopathies ^12–14^. The p38α inhibitor PF-07265803 (previously known as ARRY-797) is the only drug that progressed to the clinical trial stage for the treatment of *LMNA*-related dilated cardiomyopathy (ClinicalTrial.gov NCT03439514). Unfortunately, this program has been recently discontinued because interim results indicated it would not meet the primary endpoint. Gene therapies have also been explored to treat some laminopathies but the main focus has been on Hutchinson–Gilford progeria syndrome (HGPS). Promising results have been obtained for this LMNA-related disease using CRISPR 1.0 technology ^15,16^ and base editors ^17^, a more advanced CRISPR-based approach. For L-CMD, only one pre-clinical study reported the potential of *LMNA*-mRNA repair by spliceosome-mediated RNA trans-splicing. However, this gene therapy strategy showed a low efficiency outcome ^18^. In conclusion, there is still no promising approach for treatment of L-CMD.

Given the monogenic nature of L-CMD, this study explores the potential of a CRISPR gene-editing approach to eliminate the *Lmna* c.745C>T mutation using a mutant allele-specific RNA guide. The goal is to establish a hemizygous state for *Lmna*, aiming to reverse the pathogenic phenotype associated with L-CMD. The study involves testing this approach in different cellular models and the newly generated *Lmna^R249W^* mouse model.

## RESULTS

### Evaluation of CRISPR/Cas9 technology for the elimination of the *Lmna* c.745C>T mutation in mouse embryonic fibroblasts (MEFs): molecular characterization and phenotypic outcomes

Previous studies have demonstrated that CRISPR activity is highly dependent on the sequence complementary to the guide RNA of the CRISPR/Cas complex ^19^. Thus, alteration in just one of the twenty nucleotides of the guide results in a significant reduction in the activity of the Cas9 endonuclease on the target sequence ^20^. This cleavage specificity increases with the proximity of the point mutation to the PAM sequence ^21^. Exploiting this property of the CRISPR system, we aimed to develop a guide RNA that directs Cas9 activity preferentially to the *Lmna* c.745C>T mutant allele with minimal-to-no effect on the wild-type (WT) allele. Specifically, one guide RNA was designed to contain the cytosine-to-thymidine point mutation found at position 745 in the genomic sequence of the mutant allele (**Figure 1A**). The activity of the Cas9/sg745T complex was initially evaluated using an *in vitro* endonuclease cleavage assay with exon 4 of the *Lmna* gene, amplified from MEFs. In samples derived from MEFs harboring one or two copies of the c.745C>T mutation, two distinct bands were observed, indicating successful cleavage by the Cas9/sg745T complex. In contrast, no cleavage activity was detected in DNA amplified from *Lmna^+/+^* MEFs (**Supplemental figure 1**). Therefore, Cas9/sg745T complex activity is only observed in the presence of the mutant allele, demonstrating the specificity of the complex for the mutant allele containing the cytosine-to-thymine mutation.

**Figure 1.**
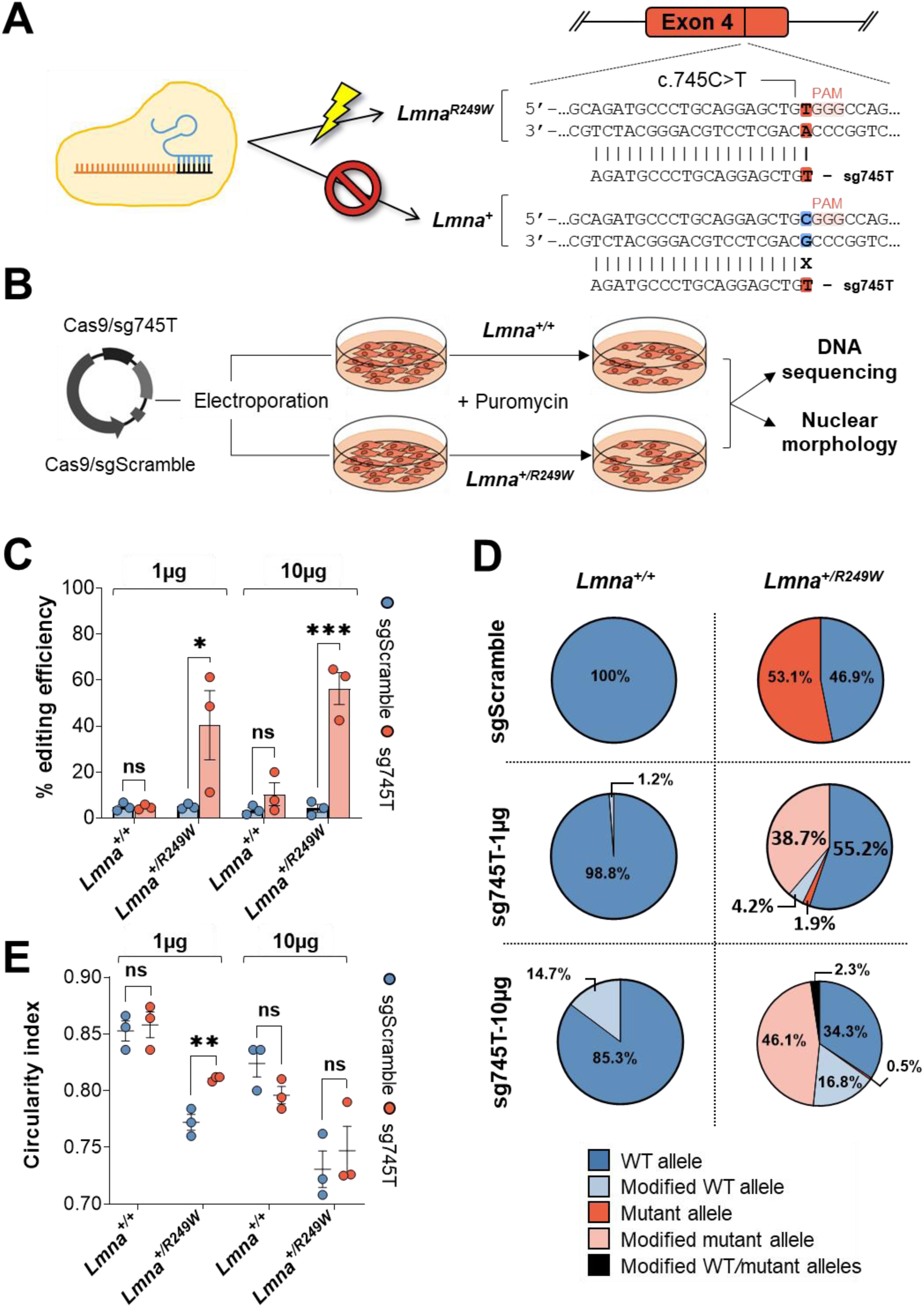
The Cas9/sg745T complex demonstrates allele-specific activity on *Lmna* c.745C>T in MEFs. **A,** Localization of the sg745T RNA guide in exon 4 of the *Lmna* gene. The nucleotide of the mutation (t=thymine, in red) is at position 745 of the coding sequence, while the nucleotide occupying the same position in the WT allele is highlighted in blue (c=cytosine). The PAM sequence is highlighted. **B,** Schematic experimental design for the assessment of Cas9/sgScramble and Cas9/sg745T complexes activity in mouse embryonic fibroblasts. Two different genotypes for the *Lmna* gene (*Lmna^+/+^* and *Lmna^+/R249W^*) were used for all experiments, and three biological replicates have been performed (n=3 pools). **C,** Analysis of CRISPR efficiency at cellular pool level using TIDE platform after nucleofection of Cas9/sgScramble and Cas9/sg745T complexes. Data are represented as mean ± SD. **D,** Percentage of reads for unmodified and modified alleles at the cellular pool level analyzed by CRISPResso2 after nucleofection with Cas9/sgScramble and Cas9/sg745T complexes. **E,** Circularity index at the cellular pool level. Data are represented as mean ± SEM. ns: non-significant differences; *: P<0.05; **: P<0.01; ***: P<0.001.

To validate our strategy, plasmids expressing Cas9 plus sg745T or a scramble RNA guide were delivered into wild-type (*Lmna^+/+^*) and heterozygous (*Lmna^+/R249W^*) MEFs carrying the c.745C>T mutation (**Figure 1B**). Two DNA amounts (1 μg and 10 μg) were tested. To determine CRISPR activity, Sanger sequencing data were analyzed using the TIDE platform (**Figure 1C**). In WT fibroblasts at low DNA concentration (1 μg), both Cas9/sgScramble (4.8 ± 1%) and Cas9/sg745T complexes (4.7 ± 0.6%) induced similar residual CRISPR activity (P=0.46). When the amount of Cas9/sg745T complexes were increased (10 μg) in *Lmna^+/+^* MEFs, this activity increased to 10.3 ± 4.9%, but there were no significant differences compared to its control Cas9/sgScramble (3.4 ± 1.3%, P=0.12). However, significant differences were detected in the heterozygous line for the c.745C>T mutation between Cas9/sg745T and Cas9/sgScramble complexes. In *Lmna^+/R249W^* MEFs, electroporation of 1 μg of Cas9/sg745T plasmid resulted in an editing efficiency significantly higher than the observed for Cas9/sgScramble (40.4% ± 15% vs 5 ± 0.7%, P=0.04). At higher concentrations (10 μg), the activity of Cas9/sg745T complexes significantly increased to 56.3 ± 6.9%, compared to the activity of 4.2 ± 1.7% detected for Cas9/sgScramble complexes (P<0.001).

These results were further confirmed through amplicon deep sequencing analysis of the target sequence using the CRISPResso2 platform (**Figure 1D**). In a *Lmna^+/+^* scenario, 100% of reads were obtained for the original WT allele in the control condition (Cas9/sgScramble). When a low concentration (1 μg) of Cas9/sg745T complexes was used, similar results were observed (98.8 ± 1.2% of reads for the original WT allele). When the concentration of Cas9/sg745T complexes was increased to 10 μg, the percentage of unmodified WT allele reads decreased to 85.3 ± 8.5%, while 14.7 ± 6% of the reads corresponded to modified WT sequences. In the case of *Lmna^+/R249W^* cells, the expression of Cas9/sgScramble complexes resulted in similar percentages of reads of WT and mutant sequences. As expected for this control, no reads for either of these two alleles were detected as modified. However, electroporation with 1 μg of Cas9/sg745T complexes led to a high percentage of reads for the modified mutant allele (38.7 ± 7.9% of total sequences). This effect was accompanied by a drastic reduction in reads for the unmodified mutant allele (1.9 ± 1% of the total). It is interesting to note that under these conditions, the percentage of modified WT allele constituted only 4.2 ± 2% of the total reads detected. When high concentrations (10 μg) of Cas9/sg745T complexes were used, the percentage of reads for the modified mutant allele increased even further to 46.1 ± 4.3% of the total. Simultaneously, the percentage of detected reads for the unmodified mutant allele was nearly zero (0.5 ± 0.5% of sequences). Interestingly, a moderated increase in modified WT sequences (16.8 ± 3.5%) was observed under these high conditions. All these analyses confirmed the specificity of Cas9/sg745T complexes for the mutant allele and demonstrated that their activity is concentration dependent.

It has been reported that nuclei of cells carrying the c.745C>T mutation exhibited compromised nuclear membrane integrity ^22^. MEFs harboring this variant show significantly lower circularity indices compared to WT cells, indicating altered nuclear morphology (**Supplemental figure 2**). The elimination of the c.745C>T mutation is presumed to restore nuclear morphology. To test this hypothesis, the circularity index of nuclei from WT and *Lmna^+/R249W^* MEFs previously transfected with Cas9/sgScramble and Cas9/sg745T complexes was quantified (**Figure 1E**). In the case of WT fibroblasts, no significant differences in circularity index were found when using Cas9/sg745T or Cas9/sgScramble complexes at any of the two concentrations used. In contrast, in a heterozygous background, an increase in the circularity of nuclei from cells nucleofected with Cas9/sg745T complexes was observed compared to those nucleofected with Cas9/sgScramble complexes. This phenotype rescue was statistically significant only in the low concentration condition (P=0.003), which, according to the deep sequencing studies, was associated with the highest percentage of unmodified WT allele (53.1 ± 3.3%) and lowest residual unmodified mutant allele (1.9 ± 1% of total reads). These results demonstrated that the specific removal of the *Lmna* c.745C>T mutation improves the nuclear morphology of heterozygous mouse embryonic fibroblasts.

### Study of specificity of CRISPR/Cas9 technology in embryos of the *Lmna^R249W^* mouse model

One-cell stage embryos were obtained from matings between *Lmna^+/+^* and *Lmna^+/R249W^* animals and nucleofected with the Cas9 and sg745T as a ribonucleoprotein complex (**Figure 2A**). Like in previous experiments, two concentrations of the Cas9:sgRNA complex were tested: a low concentration (0.61 μM) and a high concentration (8 μM). Nucleofected embryos were cultured to the blastocyst stage (E4.5 days), at which time genomic DNA was extracted, the target sequence was amplified by PCR, and editing frequency analyzed using TIDE after Sanger sequencing. A total of 130 blastocysts were analyzed. As shown in **Figure 2B**, at the low concentration (0.61 μM), only 4.5% of *Lmna^+/+^* blastocysts exhibited modifications (1 of 21), while 83.9% of *Lmna^+/R249W^* blastocysts (26 of 31) contained indels, primarily nucleotide insertions. At the high concentration (8 μM), the modification rate for *Lmna^+/R249W^* blastocysts rose to 100%, with nucleotide insertions remaining the most frequent indel type. Modifications in *Lmna^+/+^* blastocysts also increased significantly, from 4.5% to 66.7%.

**Figure 2.**
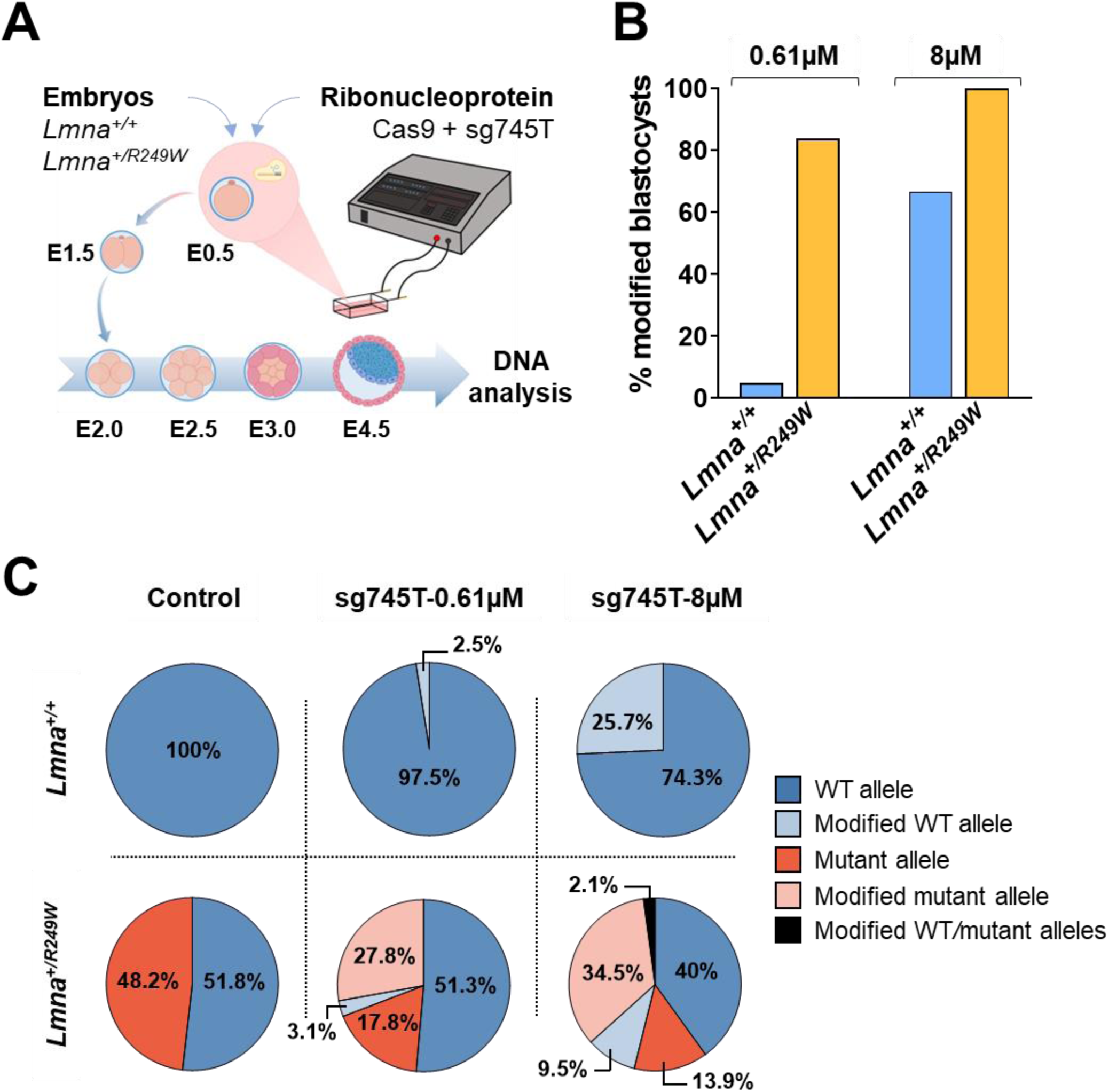
The Cas9/sg745T complex manifests high specificity for the *Lmna* c.745C>T allele in mouse embryos. **A,** Schematic experimental design for studying the activity of Cas9/sg745T complex in mouse embryos. For all experiments, two different genotypes for the *Lmna* gene (*Lmna^+/+^* and *Lmna^+/R249W^*) were used. **B,** Percentage of modified blastocysts after the introduction of Cas9/sg745T complexes. **C,** Percentage of reads for unmodified and modified alleles analyzed by CRISPResso2 in modified blastocysts after treatment with Cas9/sg745T complexes. In the control condition, 13 *Lmna^+/+^* and 17 *Lmna^+/R249W^* blastocysts were used. In the 0.61 µM condition, 21 *Lmna^+/+^* and 31 *Lmna^+/R249W^* blastocysts were employed. In the 8 µM condition, 27 *Lmna^+/+^* and 21 *Lmna^+/R249W^* blastocysts were used.

To further assess the specificity of the Cas9/sg745T complex for the c.745C>T mutant allele, blastocyst DNA was analyzed by amplicon deep sequencing (**Figure 2C**). In *Lmna^+/+^* control blastocysts (no CRISPR introduced), 100% of reads corresponded to the WT allele. At 0.61 μM, 97.5% of reads were unmodified WT, with only 2.5% showing indels. At 8 μM, unmodified WT allele reads decreased to 74.3 ± 3.9%, with modified WT reads increasing to 25.7 ± 3.8%. On the other hand, in *Lmna^+/R249W^* control blastocysts, WT and mutant alleles were equally represented (51.8 ± 2.1% vs. 48.2 ± 2.1%). At 0.61 μM, mutant allele reads decreased to 17.8 ± 1.6%, with 27.8 ± 2.5% containing Cas9-induced modifications, while WT allele modifications remained minimal (3.1 ± 1%). At 8 μM, mutant allele modifications increased to 34.5 ± 4.7%, and unmodified mutant allele reads further decreased (13.9 ± 1.9%). WT allele modifications also rose to 9.5 ± 5.6%. These results confirmed the high specificity of Cas9/sg745T complexes for the c.745C>T allele and revealed a proportional reduction in CRISPR activity targeting the mutant allele as the concentration of CRISPR complexes increased.

### Study of the potential of Cas9/sg745T mediated by AAV9 in an *in vivo*, metabolic context

After establishing the *in vitro* specificity of Cas9/sg745T complex for the mutant *Lmna* c.745C>T allele, we proceeded to validate the CRISPR-mediated strategy to eliminate the mutant allele in the *Lmna^R249W^* murine model. Our initial focus was on *Lmna^R249W/R249W^* mice, which exhibit a severe metabolic phenotype characterized by considerable growth delay, complete absence of hepatic glycogen deposits, reduced adipose tissue, and lowered body temperature. These defects culminate in premature death, with a median survival of 50 days. On the other hand, lamin A/C knockout mice are known to survive no longer than 56 days ^23^. Applying Cas9/sg745T complex therapy in the *Lmna^R249W/R249W^* model is expected to yield the elimination of the mutant allele which will generate a lamin A/C knockout mouse, which also succumbs prematurely. However, even a modest extension of survival—by up to 6 days, representing a 12% increase—would provide valuable insight into the potential therapeutic effects of this approach and allow for an initial evaluation of its impact.

To investigate the *in vivo* potential of Cas9/sg745T gene editing in this model, infections were performed in one-day-old neonates of both *Lmna^R249W/R249W^* and *Lmna^+/+^* mice using AAV9 vectors (1×10^11^ viral genomes of each vector) administered via intradermal injection in the interscapular region (**Figure 3A**). Two different viral vectors were used: one carrying the Cas9 endonuclease under the CMV promoter (AAV9-CMV-SpCas9) and the other containing the sg745T RNA guide sequence driven by the U6 promoter and the EGFP expression gene under the CMV promoter (AAV9-U6-sg745T-CMV-eGFP). The impact of this AAV-CRISPR-mediated therapy, referred to as AAV9-Cas9/sg745T or AAV-treated hereafter, on the survival and growth was evaluated compared to untreated control groups (untreated *Lmna^+/+^* and *Lmna^R249W/R249W^* mice), referred to as untreated hereafter.

**Figure 3.**
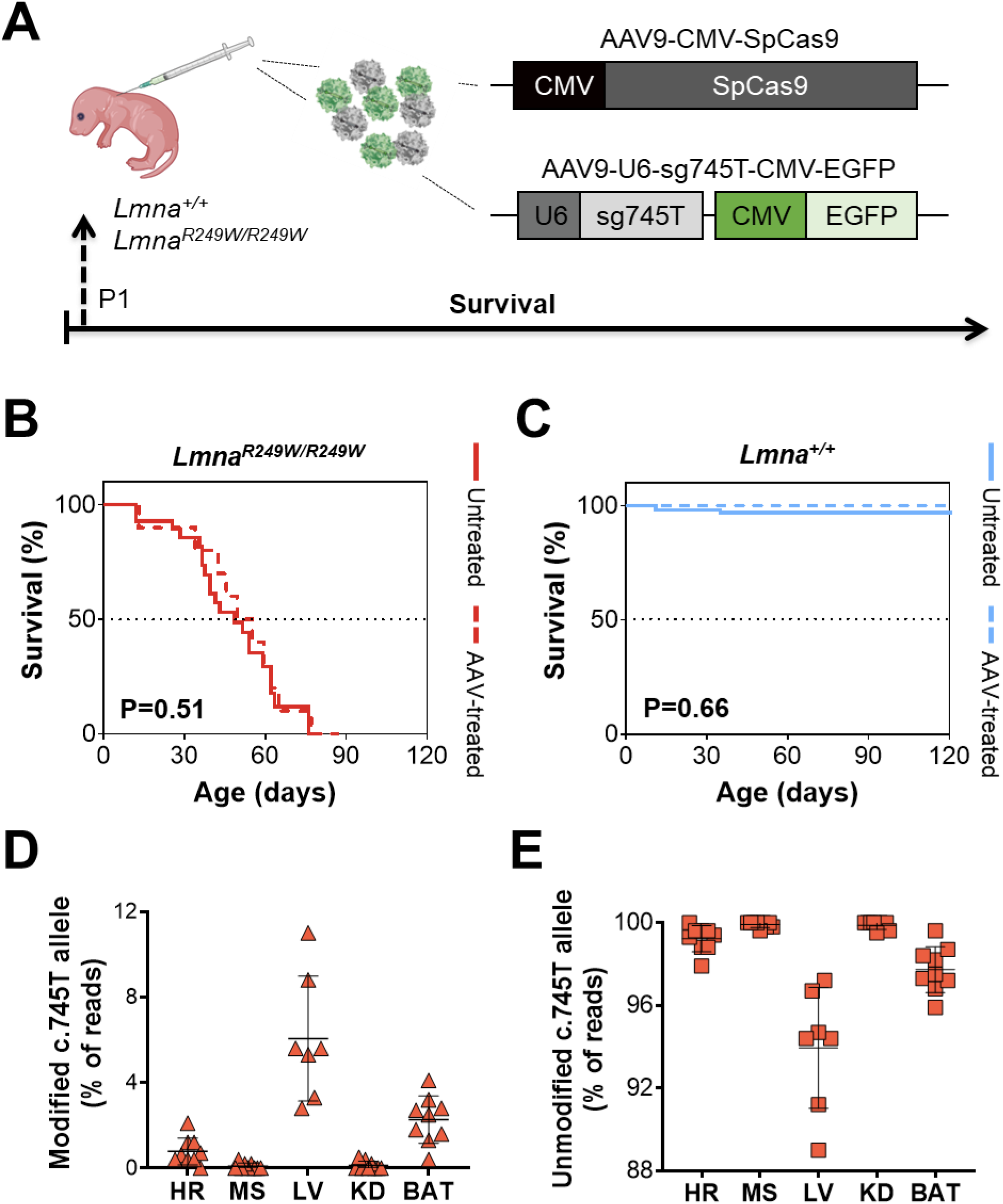
Administration of AAV9-Cas9/sg745T gene therapy does not improve survival in a homozygous background for the R249W mutation. **A,** Schematic experimental design to evaluate the survival and CRISPR activity after intradermal administration of AAV9-Cas9/sg745T treatment. AAV9-Cas/sg745T were injected into one-day-old *Lmna^R249W/R249W^* and *Lmna^+/+^* mice. Control mice received no treatment. **B,** Kaplan-Meier survival curve of untreated (n=28) and AAV-treated (n=11) *Lmna^R249W/R249W^* mice. **C,** Kaplan-Meier survival curve of untreated (n=100) and AAV-treated (n=26) *Lmna^+/+^* mice. ns: non-significant differences. **D,** Percentage of modified c.745T allele reads in *Lmna^R249W/R249W^* AAV-treated mice (n=7-9). **E**, Percentage of unmodified c.745T allele reads in *Lmna^R249W/R249W^* AAV-treated mice (n=7-9). The activity of Cas9/sg745T complex was analyzed in different tissues: heart (HR), muscle (MS), liver (LV), kidney (KD) and brown adipose tissue (BAT). Data are represented as mean ± SD.

The survival of AAV-treated *Lmna^R249W/R249W^* mice showed a modest, although not statistically significant, increase of 10% compared to untreated, control mice (55 days versus 50 days, respectively, P=0.51) (**Figure 3B**). This trend was consistent in both sexes, with AAV-treated males averaging 51 days compared with 42 days in untreated, control (P=0.29) and AAV-treated females averaging 59 days versus 55 days in untreated, control mice (P=0.89) (**Supplemental figure 3A**). In a WT background (**Figure 3C**), there was no significant survival differences between AAV-treated and untreated mice (P=0.66).

Interestingly at 21 days of age AAV-treated, *Lmna^R249W/R249W^* mice weighed significantly less than untreated controls (P=0.002) (**Supplemental figure 3B**). However, this difference was not longer apparent by 35 days of age, with AAV-treated *Lmna^R249W/R249W^* mice showing a slight, albeit non-significant increase in average body weight compared to untreated controls (P=0.20) (**Supplemental figure 3B**). When stratified by sex, the difference in body weight at 21 days of age between AAV-treated *Lmna^R249W/R249W^* and untreated control mice was only significant in males (P<0.01) (**Supplemental figure 3B**). By 35 days post-treatment, there were no significant body weight differences between treated and control mice for either sexes (**Supplemental figure 3B**). Overall, these findings indicate that AAV9-Cas9/sg745T therapy has limited impact on the survival or growth of *Lmna ^R249W/R249W^* mice.

The observed 10% survival increase in *Lmna^R249W/R249W^* mice (approaching the 56 days survival in lamin A/C knockout animals) could potentially be attributed to the editing of the two R249W alleles, leading to truncated alleles. To explore this hypothesis, CRISPR activity was analyzed in various tissues (heart, muscle, liver, kidney, and brown adipose tissue) of *Lmna^+/+^* and *Lmna^R249W/R249W^* mice aged 42-90 days post-treatment. Genomic DNA was extracted and exon 4 of the *Lmna* gene was amplified for NGS analysis. No CRISPR activity was detected in *Lmna^+/+^* mice, while modified sequences were observed in the R249W alleles of homozygous mutant mice (**Figure 3D**). Thus, a reduction in unmodified mutant alleles was observed in liver and brown adipose tissue tissues of infected *Lmna^R249W/R249W^* mice (**Figure 3E**). Despite known tropism of AAV9 tropism for cardiac and skeletal muscle, the heart exhibited only a slight reduction in the R249W allele frequency (99.2 ± 0.6%), while muscle was minimally affected (99.9 ± 0.1% R249W allele readings). The highest editing activities occurred in the liver and brown fat, where unmodified mutant allele readings decreased to 93.9 ± 2.9% and 97.7 ± 1.1%, respectively. In kidney, unmodified, allele frequencies were 99.9 ± 0.2% consistent with the serotype’s distribution. These data suggest the sg745T guide retains its specificity for the mutated R249W allele *in vivo*, although with greatly reduced activity.

Histopathological analyses of these tissues in treated *Lmna^R249W/R249W^* mice (aged 49–63 days) were compared to 35-day-old untreated controls. No significant differences or abnormalities were observed in the heart, muscle, or kidney between the groups. Although the liver exhibited the highest CRISPR activity, glycogen deposits were absent in both infected and untreated homozygous animals. In brown fat, treated mice displayed a noticeable reduction or absence of multilocular fat vacuoles, unlike the normal tissue morphology seen in untreated animals. Examination of white fat, pancreas, and spleen revealed no abnormalities in white adipose tissue or pancreas. However, lymphoid hypoplasia was detected in 50% of spleens from treated *Lmna^R249W/R249W^* mice, a feature absents in untreated controls.

In summary, AAV9-Cas9/sg745T therapy resulted in a modest, non-significant 10% increase in survival of *Lmna^R249W/R249W^* mice and showed limited CRISPR activity across tissues.

### Effect of AAV9 Cas9/sg745T gene therapy on cardiac function

*Lmna^+/R249W^* mice develop dilated cardiomyopathy, making them an ideal model to assess the impact of AAV-Cas9/sg745T gene therapy on this key pathological feature of L-CMD. *Lmna* editing efficiency was assessed in various tissues (heart, muscle, liver, kidney, and brown adipose tissue) of AAV-treated *Lmna^+/R249W^* and *Lmna^+/+^* mice. Samples were collected at 5 and 50 weeks of age to evaluate temporal changes. There was no evidence of editing in WT tissues at either time point (**Supplemental figure 4A**). Similarly, *Lmna^+/R249W^* tissues exhibited no indels in the WT allele (**Supplemental figure 4B**). However, indels were detected in the mutant allele of treated *Lmna^+/R249W^* mice (**Supplemental figure 4B**). At 50 weeks of age, the percentage of modified R249W allele reads increased significantly in the heart (from 0.4 ± 0.1% to 1.3 ± 0.3%, P=0.02) and brown adipose tissue (from 0.3 ± 0.1% to 2.0 ± 0.7%, P=0.004) from 5 to 50 weeks of age. In contrast, there was no significant difference in muscle at any time (0.07 ± 0.04% versus 0.2 ± 0.1%, P=0.23). The liver exhibited the highest allele editing activity but no significant temporal variation (3.7 ± 1.2% at 5 weeks versus 2.5 ± 1.1% at 50 weeks, P=0.30). Finally, there was minimal editing in kidney with some residual indels at 5 weeks (0.04 ± 0.04%) but not at 50 weeks of age (P=0.27).

To explore the effects of CRISPR activity, lamin A/C protein levels were evaluated in the heart, muscle, liver and brown adipose tissue of treated and untreated *Lmna^+/+^* and *Lmna^+/R249W^* mice at 50 weeks of age (**Supplemental figure 5**). Untreated *Lmna^+/R249W^* mice showed slightly reduced lamin A/C levels compared to *Lmna^+/+^* controls, with a significant reduction only in the liver (P=0.04). Treated *Lmna^+/R249W^* mice showed no significant changes in lamin A/C expression compared to untreated counterparts, consistent with the modest gene-editing levels detected in these tissues.

Despite limited editing, AAV9-Cas9/sg745T therapy significantly improved the median survival of *Lmna^+/R249W^* mice, from 437 (untreated) to 543 days (P=0.01, **Figure 4**). This 24.3% increase in lifespan was observed in males (543 vs. 415 days, P=0.01), but not in females (P=0.71) (**Supplemental figure 6**). AAV-Cas9/sg745T treatment had no impact on survival of *Lmna^+/+^* mice (**Figure 4C; Supplemental figure 6**).

**Figure 4.**
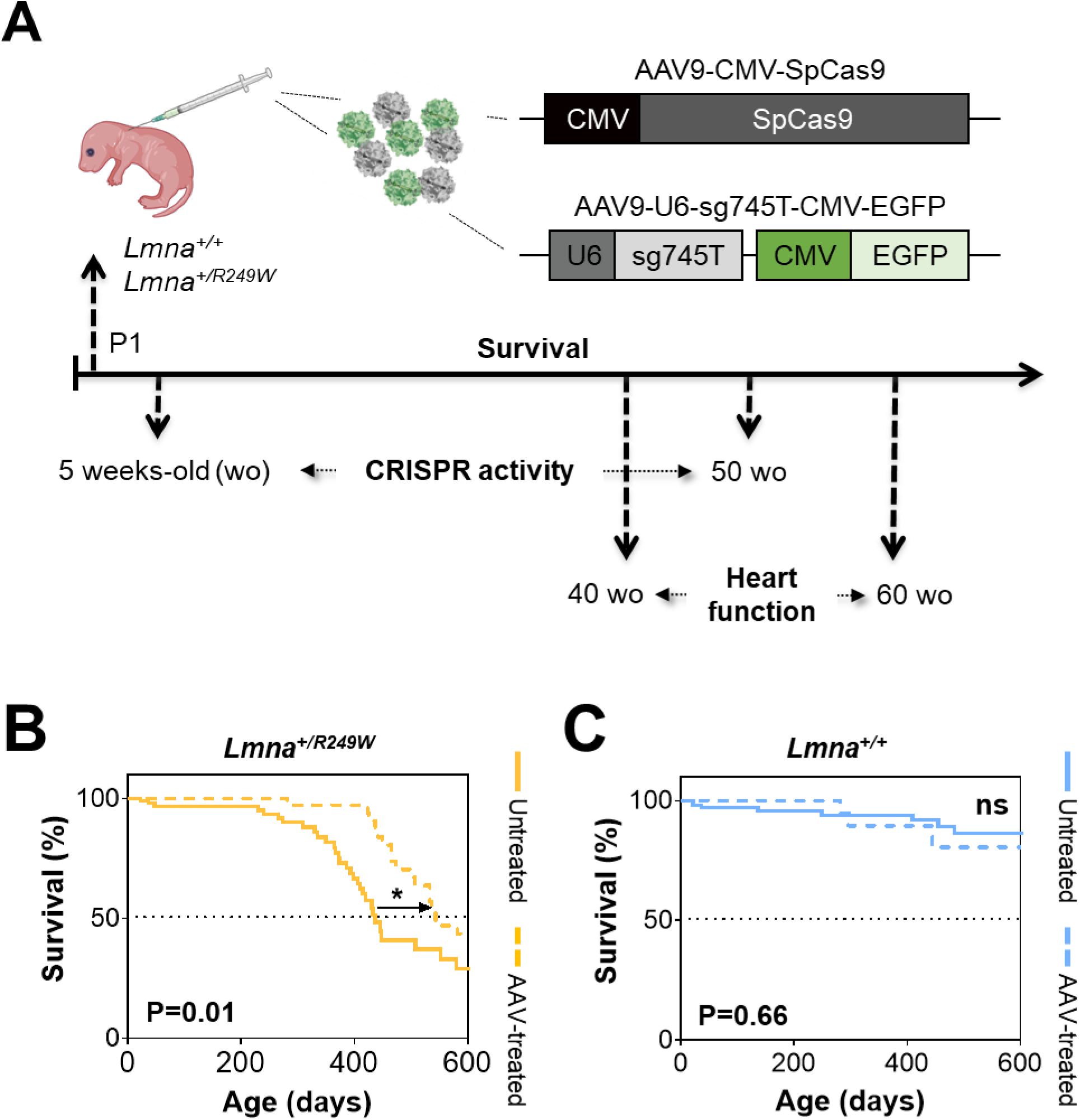
AAV9-Cas9/sg745T treatment increases survival in *Lmna^+/R249W^* mice. **A,** Schematic experimental design to evaluate the survival and cardiac function after intradermal administration of AAV9-Cas9/sg745T treatment. AAV9-Cas/sg745T were injected into one-day-old *Lmna^+/R249W^* and *Lmna^+/+^* mice. Control mice received no treatment. **B,** Kaplan-Meier survival curve of untreated (n=100) and AAV-treated (n=36) *Lmna^+/R249W^* mice. **C,** Kaplan-Meier survival curve of untreated (n=100) and AAV-treated (n=26) *Lmna^+/+^* mice. ns: non-significant differences; *: P<0.05.

Given that *Lmna^+/R249W^* mice develop dilated cardiomyopathy, the impact of AAV9-Cas9/sg745T treatment on cardiac function was assessed at 40 and 60 weeks of age using echocardiography. At 40 weeks of age, treated *Lmna^+/R249W^* mice showed significant improvement in left ventricular end-systolic (LVID, s) and end-diastolic (LVID, d) diameters (P=0.03 and P=0.03, respectively), ejection fraction (EF, P=0.02), but not for fractional shortening (FS, P=0.06) compared to untreated *Lmna^+/R249W^* mice (**Figure 5A**). These values were comparable to those of WT treated mice. At 60 weeks of age, treated *Lmna^+/R249W^* mice showed significant smaller ventricular diameters than untreated mice (LVID, s: P=0.01 and LVID, d: P=0.003, **Figure 5A**) and no significant improvement in EF and FS (P=0.07 and P=0.09, respectively, **Figure 5A**). This is consistent with progression of dilated cardiomyopathy in treated animals (LVID, s: 3.15 ± 0.88 mm, P=0.04, EF: 40.42 ± 16.81%, P=0.01).

**Figure 5.**
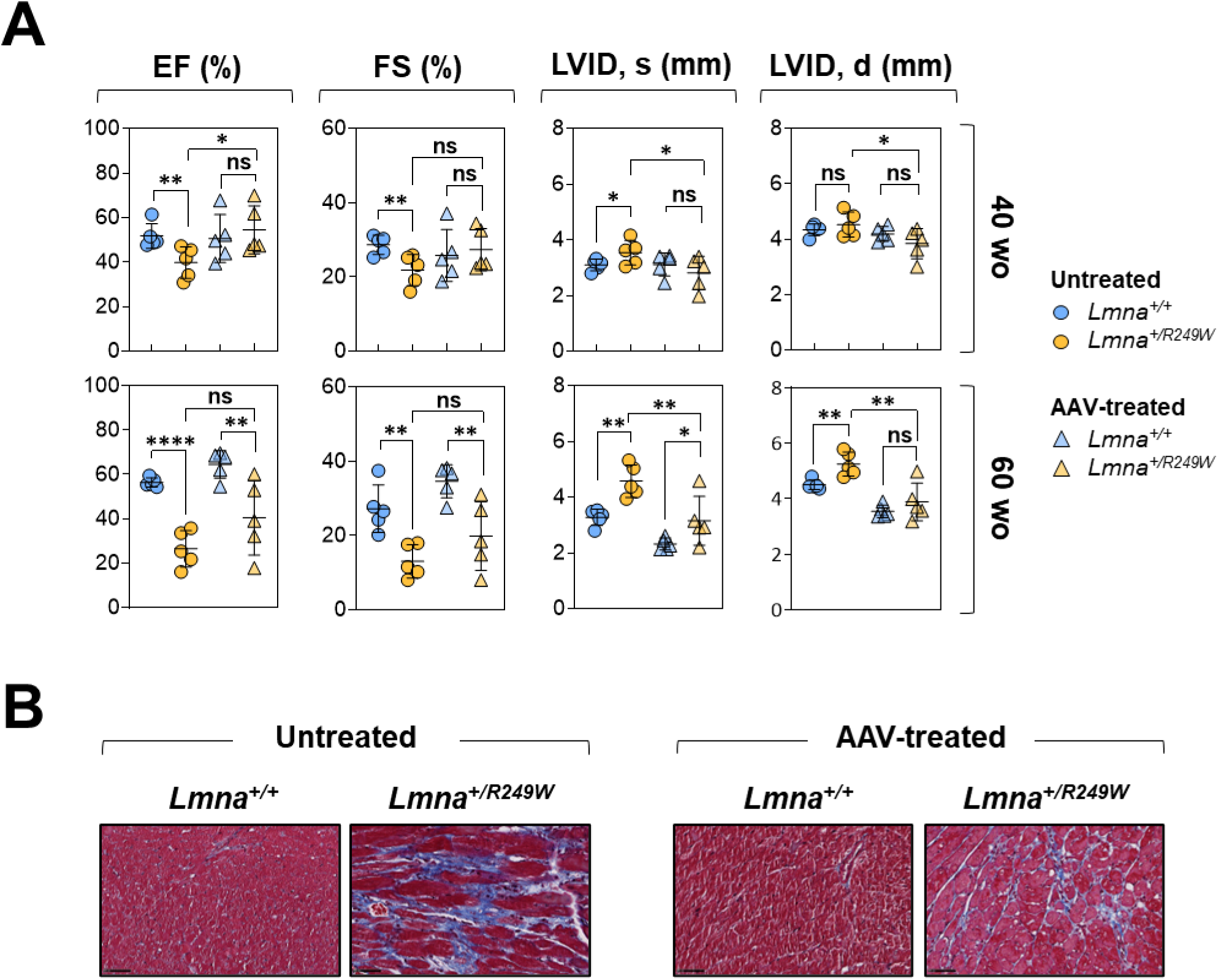
Single dose of AAV9-Cas9/sg745T treatment improves cardiac function in *Lmna^+/R249W^* mice. **A,** Echocardiographic measurements of left ventricular end-systolic internal diameter (LVID, s), left ventricular end-diastolic internal diameter (LVID, d), ejection fraction (EF) and fractional shortening (FS) at 40 and 60 weeks-old. The echocardiographic study was conducted in untreated (n=5 of each genotype) and AAV-treated (n=5 of each genotype) males. Data are represented as mean ± SD. ns: non-significant differences; *: P<0.05; **: P<0.01; ****: P<0.0001. **B,** Representative images with Masson’s trichrome staining of heart sections from untreated (n=3 of each genotype) and AAV-treated (n=3 of each genotype) males at 50 weeks of age. Scale bar: 50 µm.

In addition to the echocardiographic studies, a histopathological evaluation of the hearts from AAV9-Cas9/sg745T treated and untreated animals was conducted. This analysis revealed interstitial fibrosis in the hearts of treated *Lmna^+/R249W^* mice at 50 weeks of age, ranging from mild to severe. Severe fibrosis was observed in 33.3% of treated mice versus 100% of untreated animals (**Figure 5B**). Treated WT mice displayed no pathological changes.

In summary, a single dose of AAV9-Cas9/sg745T therapy extends the lifespan of *Lmna^+/R249W^* mice by 24.3% and partially mitigates cardiac dysfunction, as evidenced by reduced interstitial fibrosis and delayed cardiomyopathy progression. These findings highlight the potential of this gene-editing strategy for addressing *LMNA*-related cardiac diseases.

## DISCUSSION

### Efficiency and specificity of Cas9/sg745T in eliminating of the *Lmna* c.745C>T mutation

This study demonstrates the efficiency and specificity of the Cas9/sg745T complex in selectively deleting the pathogenic *Lmna* c.745C>T mutation in various cell types and a mouse model. In *Lmna^+/R249W^* fibroblasts, the Cas9/sg745T achieved editing efficiencies of 40.4% and 56.3% at plasmid concentrations of 1 and 10 μg, respectively (**Figure 1C**). In *Lmna^+/R249W^* mouse blastocysts, efficiency was even higher, reaching 83.9% and 100% at CRISPR concentrations of 0.61 and 8 μM respectively (**Figure 2B**). Although CRISPR efficiency depends on several factors, the RNA guide sequence plays a critical role in the specificity of Cas9 activity ^24,25^. It has been reported that Cas9 endonuclease may tolerate mismatches at different positions between the guide RNA and the target DNA ^26,27^. However, the seed region, consisting of the first five to eight nucleotides proximal to the PAM, is essential for initial target recognition and binding ^28–30^. Accordingly, while single nucleotide mismatches near the PAM disrupt Cas9 activity, those located in distal positions may still allow cleavage ^26,27,29^. The fact that the c.745C>T mutation resides within the seed region adjacent to the PAM may explain the high specificity of Cas9/sg745T complexes for the mutant target.

Another key aspect that impacts both the efficiency and specificity of the CRISPR complex is the dosage of Cas9 and guide RNA components. In *Lmna^+/R249W^* mouse embryonic fibroblasts low concentrations of Cas9/sg745T complexes produced a high percentage of indels in the mutant allele, with a substantial reduction in unmodified mutant alleles, leaving only 1.9% of reads unmodified (**Figure 1D**). At higher concentrations, the unmodified mutant allele was nearly eliminated, with only 0.5% of the reads remaining. A similar trend was observed in mouse embryos, where increasing concentrations of CRISPR components led to more embryos showing modifications (**Figure 2B**). In *Lmna^+/R249W^* blastocysts, the reduction of the unmodified mutant allele was less pronounced (17.8% and 13.9% of reads at low and high concentrations, respectively; **Figure 2C**). Importantly, at higher concentrations, activity was also detected in the WT allele of murine fibroblasts, with similar effects observed in embryos. This indicates that while Cas9/sg745T complexes demonstrate high specificity at lower doses, specificity decreases as concentrations increase. Previous studies suggest that high concentrations can reduce mutagenesis efficiency ^31,32^, emphasizing the importance of optimizing CRISPR component levels to balance efficiency and specificity. In our experiments, the dose-dependent nature of CRISPR activity is also evident in its impact on reducing aberrant nuclear morphology, a common cellular defect in L-CMD and other laminopathies. *Lmna^+/R249W^* fibroblasts display irregular nuclear morphology, consistent with prior reports in human and animal models ^18,33–35^. Reduction of nuclear defects has been previously demonstrated in other laminopathy models, such as *LMNA* c.1824C>T (p.G608G) in HGPS patients, where editing improved nuclear morphology ^17^. Similarly, Cas9-mediated indels targeting exon 11 of *LMNA* reduced nuclear alterations in *Lmna^G609G/G609G^* mouse fibroblasts and *Lmna^+/G608G^* patient cells ^36^.

In this study, deletion of the *Lmna* c.745C>T allele improved nuclear circularity in *Lmna^+/R249W^* fibroblasts (**Figure 1E**). However, higher CRISPR doses did not enhance phenotypic rescue, likely due to off-target activity in the WT allele. These findings highlight the importance of optimizing CRISPR dosage to balance efficiency and specificity.

### Differential effects of AAV9-Cas9/sg745T gene therapy in metabolic and cardiac contexts

This study evaluated AAV9-Cas9/sg745T therapy in metabolic and cardiac settings, revealing distinct therapeutic outcomes. In the metabolic context, treatment only provided minimal benefits in *Lmna^R249W/R249W^* animals, extending median survival by just 10% (**Figure 3B**). Editing efficiency was low across tissues, with indels detected mainly in the liver (6.1%), followed by brown adipose tissue (2.3%), heart (0.8%) and muscle (0.1%) (**Figure 3D**). These findings align with prior studies in *Lmna^G609G/G609G^* mice treated with AAV9-SaCas9, where editing was highest in the liver (13.6%) and lower in heart (5.3%) and muscle (4.1%), but resulted in a 26.4% survival increase and improved weight gain ^36^. In contrast, *Lmna^R249W/R249W^* mice showed only a slight, non-significant weight increase (**Supplemental Figure 3B**) with no restoration of hepatic glycogen stores or white fat deposits. This limited therapeutic effect could result from the low editing efficiency or the intrinsic challenge of editing both mutant alleles. Even with complete editing, elimination of both R249W alleles would result in a *Lmna* knockout, a condition associated with mortality by 56 days of age ^23^. These findings emphasize the narrow therapeutic window available in this metabolic context.

In contrast, AAV9-Cas9/sg745T therapy had a more pronounced impact in the cardiac context, improving survival and mitigating the cardiac phenotype in *Lmna^+/R249W^* mice (**Figures 4 and 5**). Similar to the homozygous setting, the highest indel levels were detected in the liver. Interestingly, indel frequencies in cardiac and brown adipose tissues increased significantly over time, from 0.4% and 0.3% at five weeks post-treatment to 1.3% and 2.0% at 50 weeks, respectively (**Supplemental Figure 4B**). These findings are consistent with previous studies targeting the *Myh6* R403Q mutation in hypertrophic cardiomyopathy models ^37^ where AAV9-SaCas9/sgRNA treatment at comparable doses resulted in low initial indel rates (0.04% and 1.19% at five and 30 weeks) in the R403Q allele that increased over time in heart. Notably, higher doses resulted in greater editing efficiencies (3%–5.9%) highlighting a dose-dependent therapeutic effect. Importantly, our study’s cardiac analysis included multiple cell populations, potentially underestimating editing efficiency. Given that cardiomyocytes—the primary targets of AAV9—constitute only 25%–35% of total cardiac cells ^38,39^, single-cell sequencing could have yield more precise measurements. Further analysis of *Lmna* expression at the mRNA level could also clarify reductions in mutant allele expression due to large deletions.

Importantly, no off-target editing was detected in the wild type allele of *Lmna^+/R249W^* mice at either five or 50 weeks post-treatment (**Supplemental figure 4B**). This aligns with findings in *Myh6^+/R403Q^* animals, where no WT allele activity was observed at 5 weeks, though higher doses resulted in of 4% and 9% loss of the WT allele by 30 weeks ^37^. In contrast, our showed no modification sin the WT allele of treated WT mice at either 5 or 50 weeks of age (**Supplemental figure 4A**) consistent with the absence of off-target effects and normal cardiac function in these animals.

While CRISPR activity did not alter LMNA protein expression levels (**Supplemental figure 5**), AAV9-Cas9/sg745T therapy significantly improved survival in *Lmna^+/R249W^* mice, extending median survival by 24.3% (**Figure 4B**). By 40 weeks post-treatment, cardiac function improved, including reduced left ventricular telesystolic and telediastolic diameters and rescued ejection fraction and fractional shortening (**Figure 5A**). However, by 60 weeks, signs of dilated cardiomyopathy emerged, though they remained less severe than in untreated heterozygous animals. Histopathological analysis confirmed these findings revealing interstitial fibrosis in treated animals, albeit to a lesser extent than in untreated mice (**Figure 5B**). These results suggest that even low levels of edited cardiac cells provide therapeutic benefits, likely by reducing the mutation burden and enhancing cellular functions. However, the lack of sustained benefits over time may reflect insufficient infection of target cells or suboptimal editing efficiency in infected cells. Comparative studies, such as those in *Myh6^+/R403Q^* mice treated with AAV9-SaCas9/sgRNA, demonstrate that medium AAV9 doses can correct cardiac hypertrophy without inducing dysfunction ^37^, supporting the use of lower viral doses to balance therapeutic efficacy and safety.

In summary, AAV9-Cas9/sg745T therapy shows promise in improving cardiac function and survival in *Lmna^+/R249W^* mice but demonstrates limited efficacy in a metabolic context. These findings underscore the need to optimize gene editing efficiency, delivery, and dosage to achieve durable therapeutic outcomes. Further investigation into cell-specific effects and contributing factors will be essential to refining this approach and ensuring its long-term efficacy.

### Gene therapy in muscular dystrophies and *LMNA*-associated diseases

CRISPR-based exon deletion therapies have been investigated in other muscular dystrophies, particularly Duchenne muscular dystrophy (DMD). Using Cas9 and RNAs guides to target and delete specific exons, these approaches have demonstrated significant therapeutic potential in preclinical studies involving murine and canine DMD models ^40–44^. Despite these advancements, the application of CRISPR/Cas9 delivered via AAVs in skeletal muscle laminopathies had not been explored prior to this study. However, similar gene editing strategies have shown promise in progeria models. For example, Beyret *et al.* delivered two RNA guides via AAV9 in transgenic *Lmna^G609G^* mice expressing Cas9, successfully reducing progerin and lamin A levels, leading to phenotypic improvements such as enhanced physical appearance, reduced weight loss, and extended survival ^15^. Similarly, Santiago-Fernández *et al.* used AAV9-Cas9 with a guide RNA targeting exon 11 of *LMNA,* resulting in reducing progerin and lamin A accumulation, improved pathological features, and increased survival rates ^36^.

Another promising therapeutic approach involves antisense oligonucleotides (ASOs), short nucleotide sequences designed to modulate mRNA expression. Given that many *LMNA* mutations exert dominant-negative effects, suppressing the mutant transcript presents a potential therapeutic strategy. Lee *et al.* demonstrated that ASO-mediated suppression of lamin A and progerin in *Lmna^G609G^* mice increased lamin C production, reduced aortic pathology, and extended lifespan ^45^. Similarly, Osorio *et al.* reported that ASO treatment reduced progerin accumulation and extended survival in progeria models ^46^. In human cells, ASOs have also been used to induce exon skipping in the *LMNA* gene ^47^. Additionally, one study has explored spliceosome-mediated RNA trans-splicing as a gene therapeutic strategy in L-CMD. In the *Lmna^ΔK^*^32^ mouse model, this technique partially corrected nuclear defects and increased *LMNA* wild-type mRNA expression in various tissues. However, the improvements were insufficient to extend lifespan ^18^.

Overall, this study represents a major advancement in gene therapy for laminopathies, successfully applying CRISPR/Cas9 to eliminate the *Lmna* c.745C>T mutation in L-CMD. It is the first potentially effective gene therapy for this rare, incurable disease and only the second CRISPR-based therapy targeting laminopathies.

These preclinical findings underscore the therapeutic potential of CRISPR/Cas9, demonstrating its ability to correct disease-specific mutations while maintaining a favorable safety profile. These results provide a foundation for further refinement and potential clinical translation of this technology for treating L-CMD and other laminopathies.

## MATERIALS AND METHODS

The details of the resources used in this research, about antibodies, cell culture media, plasmids, reagents, platforms and software, are listed in **Supplemental Table 1**.

### 1. Cell lines

All cell lines were cultured in an incubator at 37°C under an atmosphere of 5% CO_2_ and 95% humidity.

#### 1.1. Mouse embryonic fibroblasts

MEFs were isolated and immortalized from a genetically engineered mouse model constitutively expressing the *Lmna* c.745C>T, p.R249W mutation, following the standard protocol ^48^. The study utilized several MEF lines, including wild-type (*Lmna^+/+^*), heterozygous (*Lmna^+/R249W^*), and homozygous (*Lmna^R249W/R249W^*) genotypes. For genotyping and precise allele identification, the mutant *Lmna* c.745C>T allele was noted to contain an additional silent mutation (c.750T>C) and a loxP site in the intron between exons 2 and 3. Cells were cultured in Dulbecco’s Modified Eagle Medium (DMEM, 4.5 g/L glucose) supplemented with 10% fetal bovine serum and 1% penicillin-streptomycin. For nuclear morphology analysis, MEFs were plated in a 96-well glass-bottom black microplate (5,000 cells/well). After 24 hours, cells were fixed with methanol (−20°C, 5 min) and stained with Hoechst 33324 (2 μg/mL in PBS, 37°C, 15 min). Nuclear images were acquired using the Cytell Cell Imaging System, and morphology was assessed via circularity index (0–1, where 1 represents a perfect circle).

#### 1.2. Mouse embryos

Female *Lmna^+/+^* mice aged 7 to 8 weeks were hormonally stimulated to enhance follicle production. Superovulation was induced by intraperitoneal administration of equine serum gonadotropin followed 48 hours later by an injection of human chorionic gonadotropin. After the completion of the final hormonal treatment, the females were paired with *Lmna^+/R249W^* males aged 9 to 19 weeks. Successful copulation was confirmed the following morning by the presence of a vaginal plug. Subsequently, the females were euthanized by cervical dislocation, and their oviducts were carefully extracted and washed with M2 culture medium. The oviducts were then transferred to fresh M2 medium containing hyaluronidase (300 μg/mL). To retrieve embryos at E0.5 day development stage, the ampullary region of the oviducts was mechanically ruptured. The collected underwent multiple washes in M2 medium and transferred to fresh M2 medium to evaluate their viability and fertilization status. Fertilized embryos were subsequently equilibrated in KSOM medium under a layer of LiteOil Global® mineral oil to maintain optimal environmental conditions for further development.

### 2. Mouse model Lmna^R249W^

Our group previously generated a mouse model of L-CMD by introducing a heterozygous R249W mutation into the *Lmna* gene (unpublished data). The mice were maintained on a predominantly C57BL/6J background. Unless otherwise stated, both male and female mice were included in all experiments in proportional numbers.

Mice were housed in the Animal Facility of the Instituto de Salud Carlos III in ventilated polycarbonate cages, which were equipped with environmental enrichment elements. Animals had *ad libitum* access to food and water. Environmental conditions were carefully controlled, with a temperature maintained between 21–23°C, relative humidity between 55–65%, and a 12-hour light/dark cycle. For genotyping of *Lmna^R249W^* mice, genomic DNA was extracted from ear tissue by adding 500 µL of 50 mM NaOH, followed by incubation at 99-100°C until digestion was complete. Then, 100 µL of 1 M Tris-HCl (pH 7.5) was added, and the mixture was centrifuged at maximum speed for 1 minute. A region of the intron between exons 2 and 3 of *Lmna* was PCR-amplified using Genotyping-Fw and Genotyping-Rv primers (**Supplemental Table 2**). The wild-type allele produced a 217 bp band, while the allele carrying the p.R249W mutation (c.745C>T) produced a 283 bp band. All procedures involving animals were approved by the Research and Animal Welfare Ethics Committee (CEIyBA) of the Community of Madrid (PROEX164-18) and conducted in compliance with Directive 2010/63/EU on the protection of animals used for experimental and scientific purposes, as implemented in Spanish legislation through Royal Decree 53/2013.

### 3. Endonuclease cleavage assay

Genomic DNA from mouse embryonic fibroblasts was extracted using the E.Z.N.A. Tissue DNA Kit. The *Lmna* exon 4 region was amplified by PCR using 100 ng of DNA and the Lmna-Ex3-Fw and Lmna-Ex5-Rv primers (**Supplemental Table 2**). PCR products (682 bp) were confirmed by electrophoresis, excised, and purified with the E.Z.N.A. Gel Extraction Kit. DNA quantification was performed using a NanoDrop spectrophotometer (ThermoFisher Scientific, Waltham, MA, USA). For Cas9/sg745T ribonucleoprotein (RNP) complex formation, Alt-R CRISPR-Cas9 crRNA and tracrRNA (50 ng each) were incubated at a 1:1 ratio for 5 min at 95°C. The crRNA:tracrRNA complex was then mixed with Cas9 protein and 1× Cas9 buffer (5× stock: 200 mM HEPES, 1 M NaCl, 50 mM MgCl₂, 1 mM EDTA, pH 6.5) and incubated for 5 min at room temperature.

Each sample underwent two reactions: (1) a digestion reaction containing 100 ng of purified PCR product, 100 ng of Cas9 endonuclease, 100 ng of sg745T RNA guide, 2 µL of Cas9 buffer, and nuclease-free water to a final volume of 10 µL; and (2) an undigested control reaction identical to the first but without the Cas9/sg745T complex. Reactions were incubated at 37°C for 6 h, then inactivated at 65°C for 10 min. Digestion products were analyzed by electrophoresis.

### 4. Generation and introduction of CRISPR machinery into mouse cells

The RNA guide was designed to specifically target the c.745C>T mutation in exon 4 of the *Lmna* gene using the Breaking Cas Design tool (https://bioinfogp.cnb.csic.es/tools/breakingcas/) ^49^.

#### 4.1. Electroporation of mouse embryonic fibroblasts

The sgRNAs, sg745T and sgScramble (**Supplemental Table 2**), were cloned into the pSpCas9(BB)-2A-Puro vector (pX459), which contains the Cas9 endonuclease and a puromycin resistance gene. The pX459 vector was obtained from Feng Zhang’s group ^50^. A total of 1×10^6^ MEFs were nucleofected with either pX459-Cas9/sg745T or pX459-Cas9/sgScramble at low (1 μg) or high (10 μg) doses using the NEPA21 electroporator (NepaGene), following the manufacturer’s recommendations and the conditions described in **Supplemental Table 3**. After 48 hours, puromycin (2 μg/mL) was added for 3 days to enrich successfully transfected cells. The enriched pools were subsequentially expanded for further analysis.

#### 4.2. Electroporation of mouse embryos

Mouse embryos were divided into two groups: a control group (no electroporation) and an electroporated group (CRISPR machinery introduced). For nucleofection, crRNA and tracrRNA were mixed in a 1:1 ratio to form the sg745T RNA guide and pre-incubated for 5 min at 95°C. The guide was then incubated with the Cas9 protein in Opti-MEM medium for 10 min at 37°C to generate the Cas9/sg745T ribonucleoprotein complex. Embryos were nucleofected with Cas9/sg745T complexes at low (0.61 μM Cas9 endonuclease, 1.83 μM crRNA:tracrRNA) or high (8 μM Cas9 endonuclease, 24 μM crRNA:tracrRNA) doses using the NEPA21 electroporator under the conditions described in **Supplemental Table 4**. Immediately after electroporation, zygotes were transferred to KSOM medium and incubated at 37°C with 5% CO2 and 95% humidity. After 24 hours, embryos reaching the two-cell stage (E1.5 days) were selected. Both control and electroporated embryos were cultured until the blastocyst stage (E4.5 days), at which point they were collected for genomic DNA extraction and CRISPR efficiency analysis.

### 5. Genomic DNA sequencing and CRISPR activity analysis

CRISPR-induced modifications in exon 4 of the *Lmna* gene were identified using Illumina NGS and Sanger sequencing at the Genomic Unit of Instituto de Salud Carlos III. DNA was extracted from MEFs and mouse tissues using a commercial kit, while blastocyst DNA was isolated by incubating samples in 17 μL of 50 mM NaOH (95°C, 5 min), followed by neutralization with 1.7 μL of 1 M Tris (pH 8) and overnight incubation at room temperature. Illumina adapters were added to the Cas9-targeted region via PCR using DeepSeq-Fw and DeepSeq-Rv primers (**Supplemental Table 2**), followed by a second PCR to incorporate sample-specific indices. FASTQ reads were analyzed with CRISPResso2 (20 bp quantification window), identifying small insertions/deletions and quantifying regional sequences. Sanger sequencing was performed on all pools and blastocysts using DeepSeq-Fw and DeepSeq-Rv primers. Sequences were analyzed with TIDE to detect genome size modifications post-editing.

### 6. Production and administration of AAV9 viruses

AAV plasmids carrying sg745T RNA guide were constructed by VectorBuilder. Two plasmids were designed: AAV9-U6-sg745T_CMV-EGFP, containing the sg745T guide under the U6 promoter and an EGFP transgene driven by a CMV promoter, and pX551-CMV-SpCas9, expressing *Streptococcus pyogenes* Cas9 under the CMV promoter (obtained from Alex Hewitt’s group via Addgene). AAV9 vectors were produced by triple transient transfection of HEK 293 cells followed by iodixanol gradient purification as previously described ^51^. One-day-old *Lmna^+/+^*, *Lmna^+/R249W^*, and *Lmna^R249W/R249W^* mice received a single intradermal injection (1×10^11^ viral genomes per vector) in the interscapular region using 31G insulin syringes with, as previously described ^52^. Control groups remained untreated.

### 7. Phenotypic characterization of mice

#### 7.1. Body and heart weight

Body weight was recorded weekly from weaning (3 weeks old). After CO₂ euthanasia, hearts were excised and weighed using a precision balance. Heart weight was normalized to tibia length or body weight.

#### 7.2. Transthoracic echocardiography

Mice were anesthetized with inhaled isoflurane (2% induction, 1.5% maintenance) while heart rate, respiration, and temperature were monitored. Positioned supine on a heated platform, the parasternal long-axis (PLAX) view was acquired using a Vevo2100 ultrasound system (40 MHz probe, VisualSonics). Two-dimensional and M-mode images were obtained and analyzed blindly using VevoLab 5.6.1.

#### 7.3. Histological analysis

Organs (heart, gastrocnemius muscle, liver, pancreas, spleen, kidney, brown and white adipose tissue) were fixed in 4% formaldehyde, embedded in paraffin, sectioned (4–6 µm), and stained with hematoxylin-eosin. Longitudinal heart sections were also stained with Masson’s Trichrome. Images were acquired using a 3DHistech Mirax® scanner and analyzed with NDP.view2.

#### 7.4. Protein expression analysis

Tissues were frozen (−80°C), mechanically homogenized with a Precellys system (ThermoFisher) in SDS lysis buffer and incubated on ice. Lysates were centrifuged (4°C, max speed, 30 min), and protein concentration was measured via NanoDrop. Equal protein amounts were resolved on polyacrylamide gels, transferred to nitrocellulose membranes (Trans-Blot® TurboTM, Bio-Rad), and blocked in 5% milk (PBS-Tween). Membranes were incubated overnight at 4°C with primary antibodies (**Supplemental Table 1**), washed, and incubated with secondary antibodies (1 h, RT). Protein bands were detected using ECL and imaged with Amersham ImageQuantTM 800. Band intensity was quantified with ImageJ.

### 8. Statistical analysis

All analyses and graphs were generated using GraphPad Prism 8.0 software. Data are presented as mean ± SD, with “n” indicating biological replicates. Significance was set as P < 0.05. Statistical significance between two experimental groups was determined by one-tailed, unpaired Student’s t-test. Survival was analyzed using Kaplan-Meier survival curves analyzed via Mantel-Cox log-rank test.

## DATA AVAILABILITY STATEMENT

The data supporting the findings of this study are available within the article and its Supplemental Material.

## ACKNOWLEDGMENT

We thank to Ángel Zaballos, from the Genomics Unit from ISCIII, for carrying out the sequencing and the loan of indices. We want to acknowledge Raquel del Cerro, Javier Esteban and Elena Velardo, members of the Animal Facility from ISCIII, for all the help and attention dedicated to our *in vivo* experiments. We would also like to thank Patricia González, member of the Histopathology Unit from CNIO, for processing and staining of samples. We acknowledge master thesis’ student Fernando Gómez García for his support in the performance of MEFs experiments. Some of the illustrations included in the figures were created with the help of biorender.com. This research was supported by grants of Fundación Andrés Marcio, niños contra la laminopatía (TVP 259/19 to I.P.d.C.), Acción Estratégica en Salud Intramural (ISCIII, PI20CIII/00038 and PI23CIII/00041 to I.P.d.C.), Fondo Investigación Sanitaria-FIS-(PI21/00094) co-funded by the European Union, and Fundació Bosch i Aymerichand (to G.S.-B.) and Cure CMD Request for Applications (RFA), International Research Grants in Congenital Muscular Dystrophy (to I.P.d.C.).

## AUTHOR CONTRIBUTIONS

D.G.-D. acquired, analyzed and interpreted most of the data, and drafted/edited the manuscript; C.E. assisted with several *in vivo* experiments and embryos experiments; I.H. and B.V.M. performed the echocardiographies; I.H. assisted in the extraction of genomic DNA; S.C. and G.S.-B. conducted the analysis of echocardiographies; A.dM.-I. performed the histopathological study; M.S.-E. was the responsible for the production of the adeno-associated virus; I.P.d.C. developed the study concept, obtained funding, coordinated and analyzed the experimental activities and drafted/edited the manuscript. All authors have read and approved the final manuscript.

