## Supplemental Figures & Tables for "CRISPR/Cas9-mediated elimination of the *LMNA* c.745C>G pathogenic mutation enhances survival and cardiac function in *LMNA*-associated congenital muscular dystrophy"

**Supplemental figure 1. *In vitro* assessment of Cas9/sg745T complex activity in presence of the *Lmna* c.745C>T mutation.** Exon 4 of the *Lmna* gene, was amplified by PCR, resulting in a 682 bp fragment. After incubating the PCR product with and without the Cas9/sg745T complex, the reaction outcome was visualized on an agarose gel. Genomic DNA used is derived from embryonic mouse fibroblasts of three different genotype for the *Lmna* gene (WT, heterozygous and homozygous for the c.745C>T mutation). In all cases, two conditions were employed: undigested ("No C/sg": not incubated with Cas9 and sg745T) and digested ("Cas/sg745T": incubated with Cas9 and sg745T guide). The white arrow indicates the band from the undigested PCR product (size of 682 bp), while black arrows correspond to the digestion result (two bands with sizes of 394 and 288 bp).

**Supplemental figure 2. The nuclear morphology of MEFs containing the *Lmna* c.745C>T mutation is significantly altered.** Circularity index of cell lines with three different *Lmna* genotypes (WT, heterozygous, and homozygous for the c.745C>T mutation). Data are represented as mean ± SD, with results obtained from five technical replicates (n=5). ****: P<0.0001.

**Supplemental figure 3. Administration of AAV9-Cas9/sg745T gene therapy does not improve survival or body weight in both *Lmna*$^{R249W/R249W}$ males and females. A,** Kaplan-Meier survival curve of males (upper graph) (untreated: n=50 for *Lmna*$^{+/+}$ and n=14 for *Lmna*$^{R249W/R249W}$; and AAV-treated: n=13 for *Lmna*$^{+/+}$ and n=11 for *Lmna*$^{R249W/R249W}$) and females (bottom graph) (untreated: n=50 for *Lmna*$^{+/+}$ and n=8 for *Lmna*$^{R249W/R249W}$; and AAV-treated: n=13 for *Lmna*$^{+/+}$ and n=3 for *Lmna*$^{R249W/R249W}$). **B,** Comparison of body weight in untreated and AAV-treated *Lmna*$^{R249W/R249W}$ males (upper graph) and females (bottom graph) at 21 and 35 days of age. Data are presented as mean values ± SD. ns: non-significant differences, **: P<0.01.

**Supplemental figure 4. Quantification of indel generation in *Lmna*$^{+/+}$ and *Lmna*$^{+/R249W}$ mice infected with AAV9-Cas9/sg745T. A,** Percentage of modified WT allele reads in *Lmna*$^{+/+}$ AAV-treated mice at 5 (n=5) and 50 (n=3) weeks-old. **B,** Percentage of modified WT (upper graph) and modified c.745T (bottom graph) alleles reads in *Lmna*$^{+/R249W}$ AAV-treated mice at 5 (n=7) and 50 (n=3) weeks of age. The activity of the Cas9/sg745T complex was analyzed in different tissues: heart (HR), muscle (MS), liver (LV), kidney (KD) and brown adipose tissue (BAT). Data are presented as mean values ± SD. ns: non-significant differences; *: P<0.05; **: P<0.01.

**Supplemental figure 5. *Lmna*$^{+/R249W}$ mice treated with AAV9-Cas9/sg745T show no differences in lamin A/C expression levels compared to untreated ones. A,** Representative Western blot of lamin A/C protein expression from lysates obtained from heart (HR), muscle (MS), liver (LV) and brown adipose tissue (BAT). GAPDH was used as a loading control. **B,** Relative quantification of total lamin A/C normalized to GAPDH in heart, muscle, liver, and brown adipose tissue. Data are presented mean values ± SD. ns: non-significant differences, *: P<0.05. Samples from 3 mice aged 50 weeks were used for each of the compared groups.

**Supplemental figure 6. Administration of AAV9-Cas9/sg745T gene therapy increases survival in *Lmna*$^{+/R249W}$ males.** Kaplan-Meier survival curve of males (untreated: n=50 for *Lmna*$^{+/+}$ and n=55 for *Lmna*$^{+/R249W}$; and AAV-treated: n=13 for *Lmna*$^{+/+}$ and n=16 for *Lmna*$^{+/R249W}$) and females (untreated: n=50 for *Lmna*$^{+/+}$ and n=45 for *Lmna*$^{+/R249W}$; and AAV-treated: n=13 for *Lmna*$^{+/+}$ and n=20 for *Lmna*$^{+/R249W}$).

### SUPP. FIGURE 1

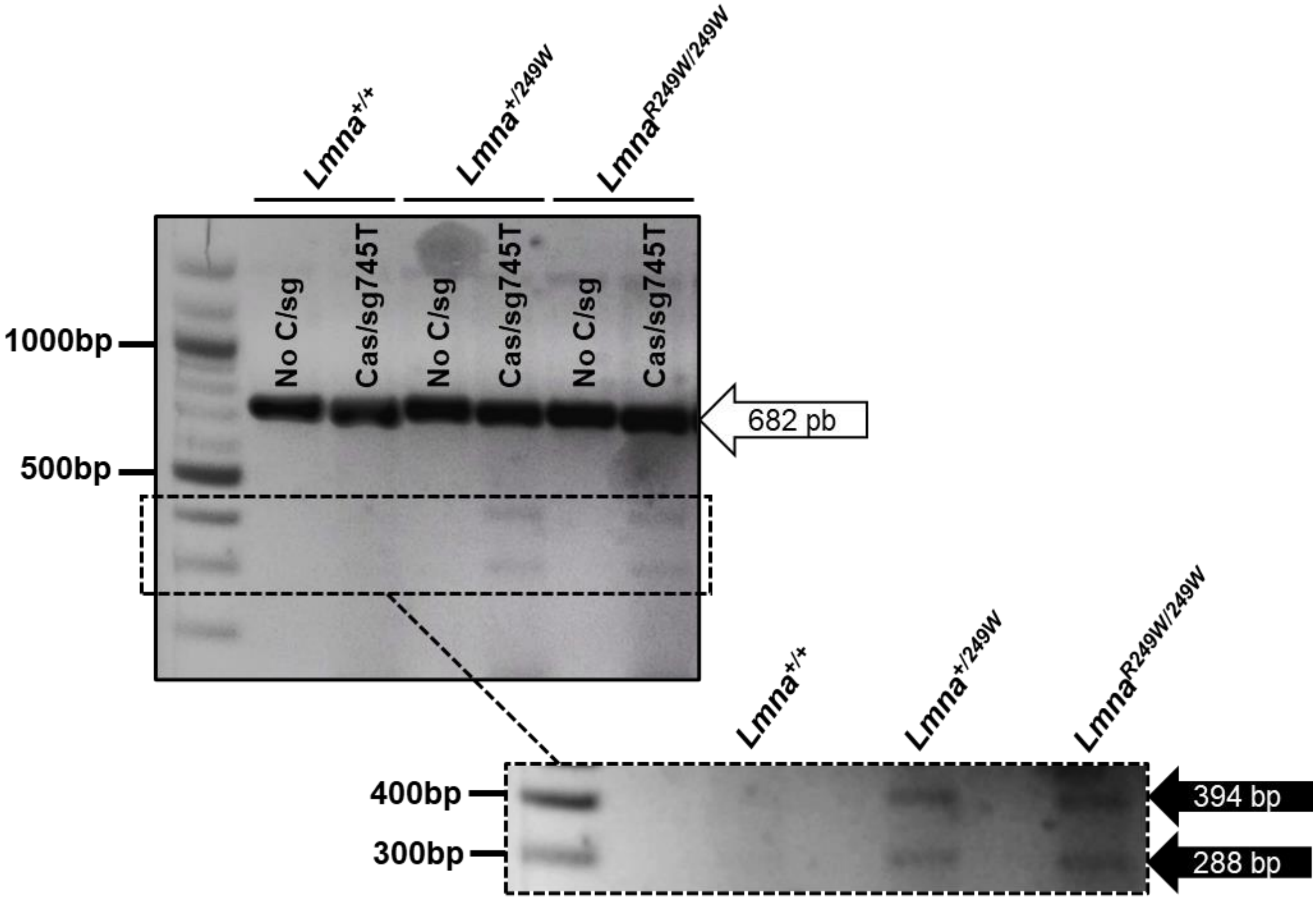

### SUPP. FIGURE 2

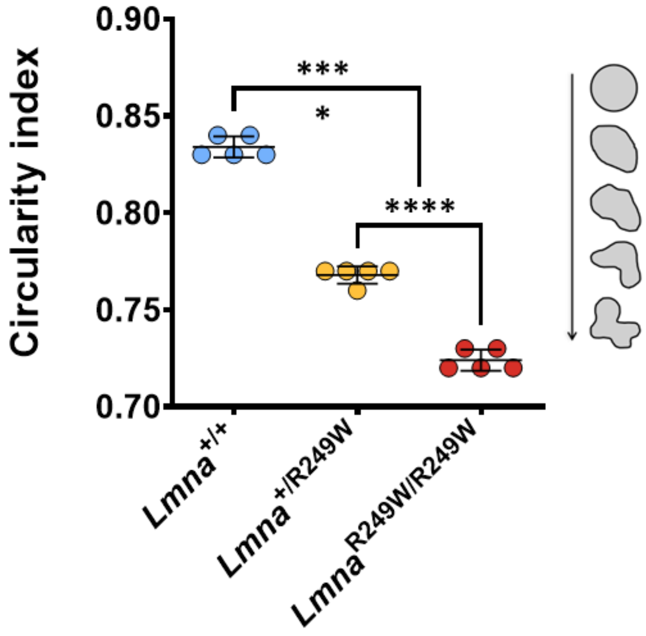

### SUPP. FIGURE 3

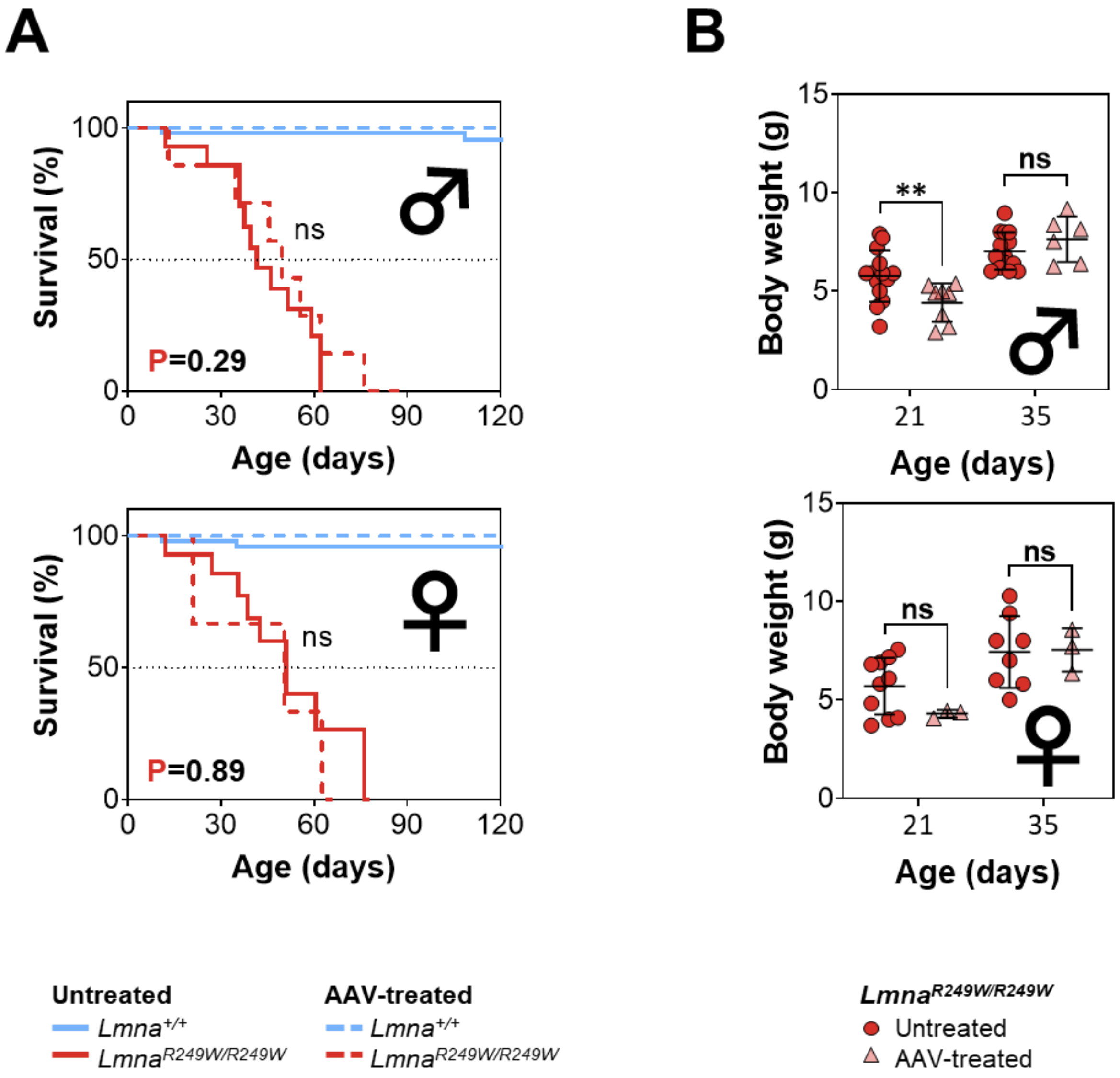

### SUPP. FIGURE 4

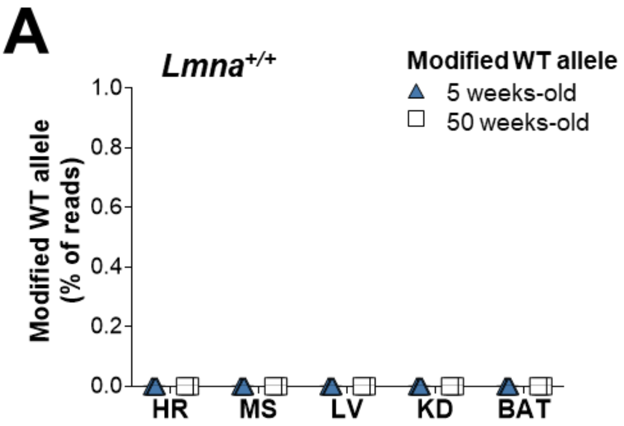


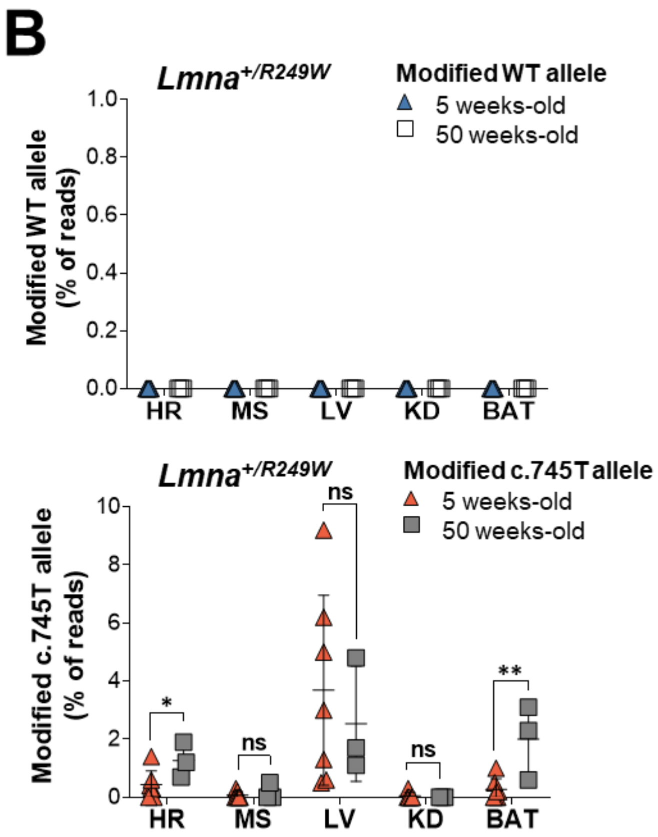

### SUPP. FIGURE 5

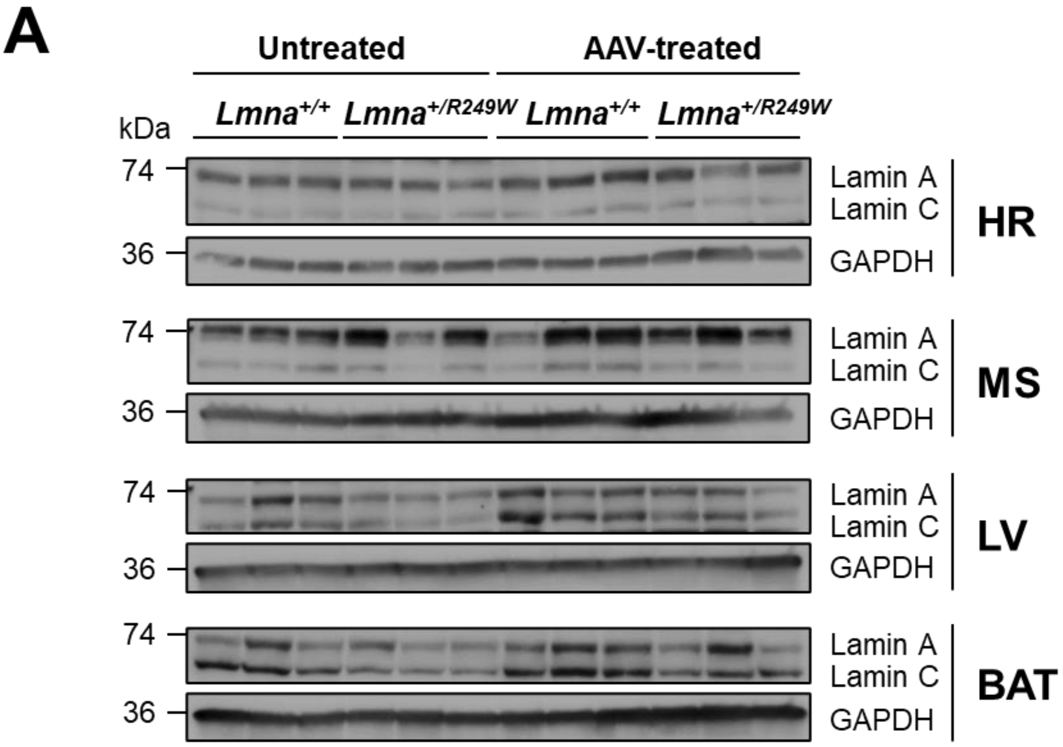


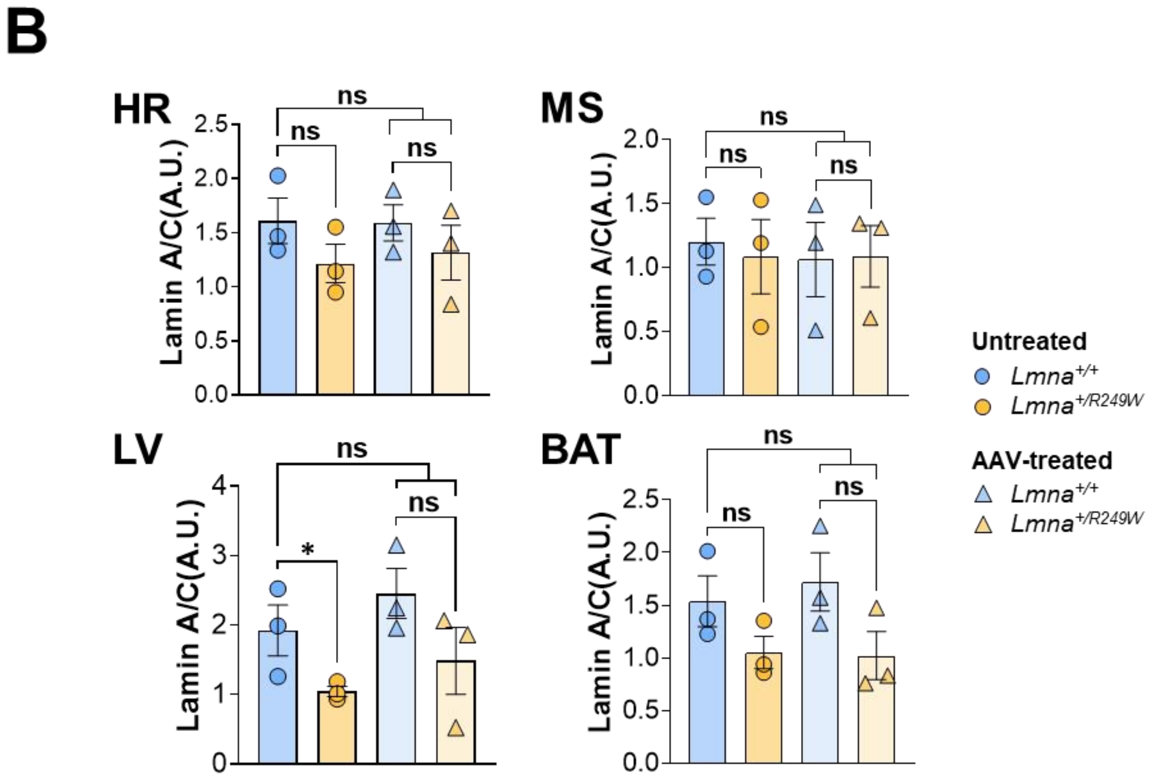

### SUPP. FIGURE 6

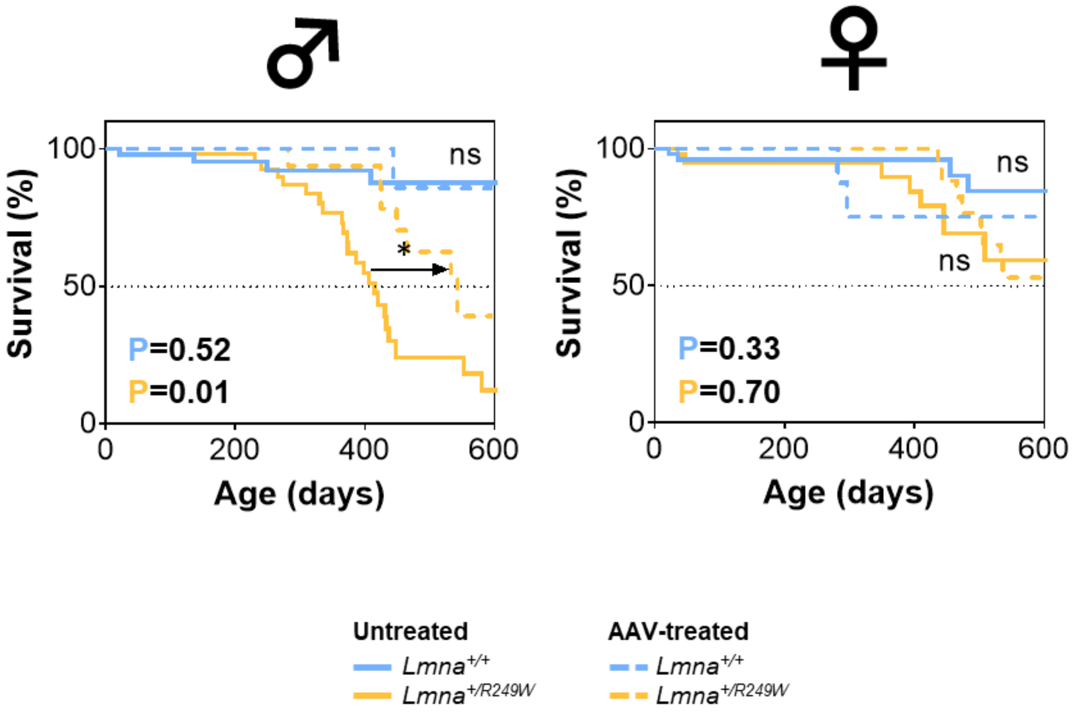

### SUPPLEMENTAL TABLES

**Supplemental Table 1. Resources used in this work.**

| Antibodies (working dilution used) |
|---|
| GAPDH (14C10) rabbit mAb (1:200) / Cell Signaling (Danvers, MA, USA) (#2118) |
| HRP-labelled anti-rabbit secondary antibody (1:5000) / GE Healthcare (Chicago, IL, USA) (NA934-1mL) |
| Lamin A/C rabbit polyclonal antibody (1:2000) / Proteintech (Manchester, UK) (10298-1-AP) |
| **Cell culture media** |
| DMEM (Dulbecco's modified Eagle's medium) high glucose / Invitrogen (Waltham, MA, USA) (61965-026) |
| Fetal bovine serum / Sigma-Aldrich (St. Louis, MI, USA) (#F7524-500mL) |
| KSOM medium / Sigma (St. Louis, MI, USA) (MR-101-D) |
| LiteOil Global® mineral oil / DiviLab (Buenos aires, Argentina) (LGOL-100, 100mL) |
| M2 medium / Sigma-Aldrich (St. Louis, MI, USA) (M7167-100mL) |
| Penicillin/streptomycin / Lonza (Basel, Switzerland) (#DE17-602E) |
| **Plasmids** |
| pX459 vector (pSpCas9(BB)-2A-Puro) / Addgene (Watertown, MA, USA) (#62988) |
| pX551-CMV-SpCas9 vector / Addgene (Watertown, MA, USA) (#107024) |
| U6-sg745T_CMV-EGFP vector / VectorBuilder (Chicago, IL, USA) |
| **Reagents** |
| Alt-R® CRISPR-Cas9 crRNA, 10 nmol / IDT (Newark, NJ, USA) |
| Alt-R® CRISPR-Cas9 tracrRNA, ATTO™ 550, 20 nmol / IDT (Newark, NJ, USA) (1075928) |
| BD Safet-GlideTM Insulin, 3/10mL 31G x 5/16 TW / Becton Dickinson (Flankin Lakes, NJ, USA) (305937) |
| Cell culture microplate 96 wells / Greiner Bio-one (Kremsmünster, Austria) (655986) |
| Criterion TGX Stain-Free Precast Gels / Bio-Rad (Hercules, CA, USA) (#5678084) |
| ECL western blotting system / ThermoFisher Scientific (Waltham, MA, USA) (32106) |
| EDTA Disodium salt dihydrate / Panreac (Castellar del Vallès, Spain) (131669.1210) |
| Equine serum gonadotropin hormone / Sigma (St. Louis, MI, USA) (G4877, 9002-70-4) |
| Formaldehyde / VWR BDH Chemicals (Mumbai, India) (11699408) |
| GenCrispr Cas9 nuclease / GenScript (Piscataway, NJ, USA) (Z03393) |
| Hoechst 33324 / ThermoFisher Scientific (Waltham, MA, USA) (H3570) |
| Human chorionic gonadotropin hormone / Sigma (St. Louis, MI, USA) (CG-10, 9002-61-3) |
| Hyaluronidase / StemCell (Vancouver, BC, Canada) (#07461) |
| Indices / Nextera XT index kit / Illumina (San Diego, CA, USA) |
| ISOFLUTEK® 1000 mg/g / Laboratorios Karizoo (Caldes de Montbui, Spain) (100603) |
| Methanol / Panreac AppliChem (Castellar del Vallès, Spain) (#131091.1612) |
| NaOH / Panreac (Castellar del Vallès, Spain) (131659) |
| Nuclease Free Water / IDT (Newark, NJ, USA) (10-04-02-01) |
| Opti-MEM™ reduced serum medium / Gibco (Waltham, MA, USA) (31985-070) |
| PBS (phosphate-buffered saline) / Lonza (Basel, Switzerland) (#BE17-515Q) |
| Puromycin / InvivoGen (San Diego, CA, USA) (ant-pr-1) |
| SDS / Sigma (St. Louis, MI, USA) (L3771-100G) |
| Trans-Blot Turbo Midi 0.2 µm Nitrocellulose Transfer Packs / Bio-Rad (Hercules, CA, USA) (#1704159) |
| Tris / Panreac (Castellar del Vallès, Spain) (A1086,1000) |
| Tween-20 / Fisher Bioreagents (Waltham, MA, USA) (BP337-100) |

| Comercial kits |
|---|
| E.Z.N.A.® Tissue DNA Kit (V-Spin) / Omega Bio-Tek (Norcross, GA, USA) (D03396-02) |
| E.Z.N.A.® Gel Extraction kit (V-Spin column) / Omega Bio-Tek (Norcross, GA, USA) (D2500-02) |
| **Platforms and softwares** |
| Breaking Cas Design (https://bioinfogp.cnb.csic.es/tools/breakingcas/) |
| CRISPResso2 (http://crispresso.pinellolab.partners.org/) |
| ImageJ (US National Institutes of Health, Bethesda, MD, USA) |
| NDP.view2 (Hamamatsu) |
| Prism 8 (GrahpPad Sowtware, Inc, San Diego, CA, USA) |
| TIDE (https://tide.nki.nl/) |
| Vevo LAB (FUJIFILM VisualSonics, Inc) |

**Supplemental Table 2. Primer and RNA guides sequences used in this work.** *

| Primer name and 5' to 3' sequence |
|---|
| Lmna-Ex3-Fw: CTGGGAGAGGCTAAGAAGCA |
| Lmna-Ex5-Rv: GTCAATGCGGATTCGAGACT |
| Genotyping-Fw: CTTCTGCCATGTAGGCTCTAAG |
| Genotyping-Rv: ATGCCAAAGGAGAGGTGATG |
| DeepSeq-Fw: tcgtcggcagcgtcagatgtgtataagagacagGCTTCTAAGGAACCATTGCGA |
| DeepSeq-Rv: gtctcgtgggctcggagatgtgtataagagacagACCAGGGAGAGGACAGGAT |
| **RNA guide name and 5' to 3' sequence** |
| sg745T: AGATGCCCTGCAGGAGCTGT |
| sgScramble: GTGTAGTTCGACCATTCGTG |

*Additional sequences added to the primer sequence for Deep sequencing PCR are indicated in lower case.

**Supplemental Table 3. Nucleofection conditions for mouse embryonic fibroblasts using NEPA21 electroporator.**

| Poring pulse | | | | | |
|---|---|---|---|---|---|
| Voltage | Lenght (ms) | Interval (ms) | No. | D. Rate (%) | Polarity |
| 200 | 5 | 50 | 2 | 10 | + |
| **Transfer pulse** | | | | | |
| Voltage | Lenght (ms) | Interval (ms) | No. | D. Rate (%) | Polarity |
| 20 | 50 | 50 | 5 | 40 | +/- |

**Supplemental Table 4. Nucleofection conditions for mouse embryos using NEPA21 electroporator.**

| Poring pulse | | | | | |
|---|---|---|---|---|---|
| Voltage | Lenght (ms) | Interval (ms) | No. | D. Rate (%) | Polarity |
| 225 | 1.5 | 50 | 4 | 10 | + |
| **Transfer pulse** | | | | | |
| Voltage | Lenght (ms) | Interval (ms) | No. | D. Rate (%) | Polarity |
| 20 | 50 | 50 | 5 | 40 | +/- |
